# Machine learning guided association of adverse drug reactions with *in vitro* target-based pharmacology

**DOI:** 10.1101/750950

**Authors:** Robert Ietswaart, Seda Arat, Amanda X. Chen, Saman Farahmand, Bumjun Kim, William DuMouchel, Duncan Armstrong, Alexander Fekete, Jeffrey J. Sutherland, Laszlo Urban

## Abstract

Adverse drug reactions (ADRs) are one of the leading causes of morbidity and mortality in health care. Understanding which drug targets are linked to ADRs can lead to the development of safer medicines. Here, we analyze *in vitro* secondary pharmacology of common (off) targets for 2134 marketed drugs. To associate these drugs with human ADRs, we utilized FDA Adverse Event Reports and developed random forest models that predict ADR occurrences from *in vitro* pharmacological profiles. By evaluating Gini importance scores of model features, we identify 221 target-ADR associations, which co-occur in PubMed abstracts to a greater extent than expected by chance. Among these are established relations, such as the association of *in vitro* hERG binding with cardiac arrhythmias, which further validate our machine learning approach. Evidence on bile acid metabolism supports our identification of associations between the Bile Salt Export Pump and renal, thyroid, lipid metabolism, respiratory tract and central nervous system disorders. Unexpectedly, our model suggests PDE3 is associated with 40 ADRs. These associations provide a comprehensive resource to support drug development and human biology studies.

---

Toxicity is one of the major causes of termination, withdrawal, or labeling of a drug candidate or drug, other than lack of efficacy ^1–3^. There is an urgent need to better identify toxic on- and off-target effects on vital organ systems especially for cardiovascular, renal, hepatic and central nervous system (CNS)-related toxicities; furthermore, there is a desire to reduce cost and labor in preclinical assays and drug testing on non-human species ^4–6^. *In vitro* pharmacological assays have been widely used to screen for possible off-targets and potential adverse effects and eliminate compounds that are not safe enough in the drug development stage as early as possible ^5,7^. However, systematic prediction of compound safety and potential adverse events associated with a compound is still a challenge for the pharmaceutical industry.

Machine learning can be very insightful for many different stages of drug discovery and development, such as automation in pharmacology assays, clinical trials, and basic science research. Previous studies have focused on predicting structure-function relationships based on chemical structure of small molecules and potency assays that probe the physicochemical properties of compounds to estimate associations with off-targets ^8^. However, the diversity of structures that interact with targets, even when they are well described like human Ether-a-go-go-related gene (hERG), make it challenging to produce reliable models ^9^. Several papers provide small, hand-curated databases providing up to 70 pharmacological targets (i.e. receptors, ion channels, transporters, etc.) with established links to adverse side effects based on a scientific literature search ^5,7,10,11^. Mirams et al. recently described how integration of data from multiple ion channels (e.g. hERG, sodium, L-type calcium) provided improved *in silico* prediction of torsadogenic risk ^12^. Chen et al. proposed a machine learning approach to predict adverse drug reaction (ADR) outcomes for given patient characteristics and drug usage ^13^. Another study highlights the importance of predicting the likelihood of clinical trial side effects using human genetic studies of drug-targeted proteins ^14^. From a pharmacogenomics perspective, predicting drug-target interactions using pharmacological similarities of drugs and the US Food and Drug Administration (FDA) Adverse Event Reporting System (FAERS ^15^) can be beneficial for drug repositioning and repurposing ^16^.

FAERS is a voluntary, post-marketing pharmacovigilance tool that can be used to monitor the clinical performance of drugs. In this study, we explore an alternative use of FAERS data to predict compound safety using Medical Dictionary for Regulatory Activities (MedDRA® ^17^) terms, which we envision to be useful for future preclinical studies. Our machine learning approach is different from the aforementioned approaches because we not only predict adverse drug reaction occurrences of drugs but most importantly also extract biologically meaningful target-ADR links. Using an *in vitro* secondary pharmacology dataset of more than 2,000 marketed or withdrawn drugs (see Methods), we built a random forest model to predict drug-ADR and target-ADR associations. We validate drug-ADR predictions through systematic Side Effect Resource (SIDER) drug label analysis and 221 target-ADR predictions through systematic literature co-occurrence analysis. Furthermore, we find canonical target-ADR associations, such as hERG binding causing cardiac arrhythmias. We also encountered unexpected associations which warrant further investigations, such as a link between Phosphodiesterase 3 (PDE3) and several ADRs, including congenital renal and urinary tract disorders. We conclude our study with potential targets that are associated with cardiovascular and renal ADRs to demonstrate the utility and possible impact of this method in drug development and preclinical safety sciences by enabling prediction of ADRs from *in vitro* pharmacological profiles.

## Results

### Systematic *in vitro* pharmacology of marketed and withdrawn drugs

To link gene targets to ADR occurrence, we utilized *in vitro* pharmacology assay data for 2134 marketed or withdrawn drugs, generated by Novartis, and ADR reports from FAERS (Figure 1A, Supplementary Table 1). Withdrawn drugs and their assay data are also included due to the fact that they are associated with a plethora of ADRs, and thereby constitute an important resource for our predictive approach. Figure 1B summarizes the top 50% of frequently occurring primary indications, classified by the Anatomical Therapeutic Chemical (ATC) codes, of the 2134 drugs using a word cloud. The categories that have the highest number of compounds are antibacterial, ophthalmological, and antineoplastic drugs. The *in vitro* pharmacology assay data includes AC_50_ values for each drug at up to 218 different assays for 184 gene targets (see Supplementary Table 2 for a list of target assays). There are 6 classes of these 184 gene targets, with the majority (47%) of targets falling into G protein-coupled receptors (GPCRs) (Figure 1C), which is a dominant, widely studied drug target family, broadly represented by marketed drugs ^18^. Figure 1D is a heatmap visualization of the *in vitro* pharmacology assay data, where each row is a drug, grouped by their ATC anatomical main group terms ^19^; each column is a target assay, grouped by target class; and each value is the AC_50_ of drugs for target assays. Even though 70% of drug-assay combinations have not been tested, i.e. these combinations have NA value for AC_50_, our data indicate relatively uniform assaying with respect to the different drug classes.

**Figure 1.**
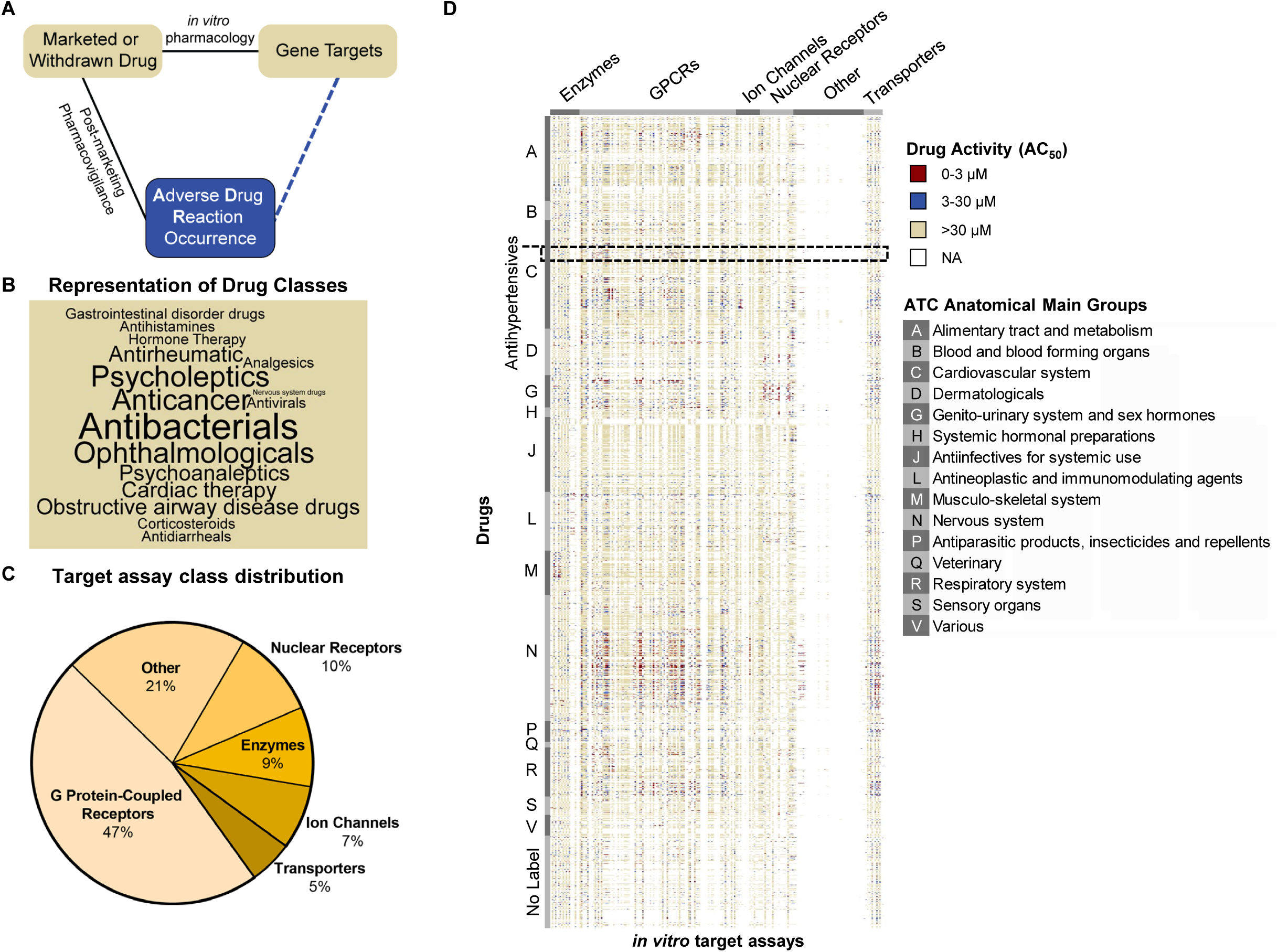
Major elements of the target-ADR association analysis. A. **Schematic outline of target-ADR pair determinations.** The observed relations (solid lines) between drugs and adverse drug reactions (ADRs) are determined by post-marketing pharmacovigilance and between drugs and their (off) targets by *in vitro* pharmacology. This approach enables prediction of associations (dashed line) between targets and ADRs through random forest modeling. B. **Representation of drug classes in word cloud.** The cloud displays the top 50% most frequently occurring drug classes, representing 2134 drugs, in the Novartis *in vitro* pharmacology data warehouse. Size of the font of the drug class reflects the number of associated drugs. C. **Target class distribution in the Novartis *in vitro* secondary pharmacology assay panel.** The 184 targets in the Novartis assay panel cover 6 target classes. Almost half of the target assays belong to the G protein-coupled receptor (GPCR) class. D. **Novartis target panel potency (AC**_**50**_**) heatmap.** The profile consists of the AC_50_ values of 184 target assays for 2134 drugs. We considered an AC_50_ value less than 3 μM as highly active (red), between 3 μM and 30 μM as active (blue), and greater than 30 μM as inactive (yellow). No data for a drug-target pair is labeled as NA (white). Drugs are grouped (vertically) by their Anatomical Therapeutic Chemical (ATC) codes. Assays are grouped (horizontally) by target class.

### Analysis of adverse event reports from FAERS connects drugs with human ADRs

We queried FAERS ^15^ using openFDA ^20^ for 2134 marketed or withdrawn drugs in October 2018 (FAERS Q4_2018 version; covering all reports from January 2004 to October 2018) and retrieved 671,143 adverse event reports using our data extraction criteria (Figure 2A). We only included reports which were submitted by physicians and were annotated as the primary suspect drug ^21^. There are 464 drugs that did not have a matching name in FAERS, 341 drugs that did not have any adverse event reports, and 1329 drugs with at least 1 adverse event report. We developed a significance test based on a binomial null distribution and false discovery rate (FDR) multiple testing correction to determine if the observed ADR occurrence was significantly high to be classified as an association (or alternatively no association) between ADR and drug (see Methods for detail). The resulting drug-ADR associations corresponded strongly (odds ratio = 11, χ^2^-test, p-value < 10^−16^) with those identified with ERAM (Empirical-Bayes Regression-adjusted Arithmetic Mean), an established Bayesian method based on the proportional reporting ratio adjusted for covariates (age group, sex and reporting year) and concomitant drugs ^22–25^. Overall, we observe a positive trend between the number of adverse event reports and the number of ADR associations (Figure 2B). Antineoplastic and immunomodulatory drugs (Figure 2B, blue, N=155) have many ADR associations while the extent of ADR association for antihypertensive drugs (Figure 2B, red, N=35) varies more widely. As an example, we visualized our drug-ADR associations (Figure 2C), in which ADRs are grouped by MedDRA System Organ Class (SOC) level terms and drugs are grouped by ATC anatomical main group terms ^19^, revealing that ADRs are widespread across organs caused by antineoplastic and immunomodulating agents (Figure 2C, label L), as well as nervous system drugs (Figure 2C, label N).

**Figure 2.**
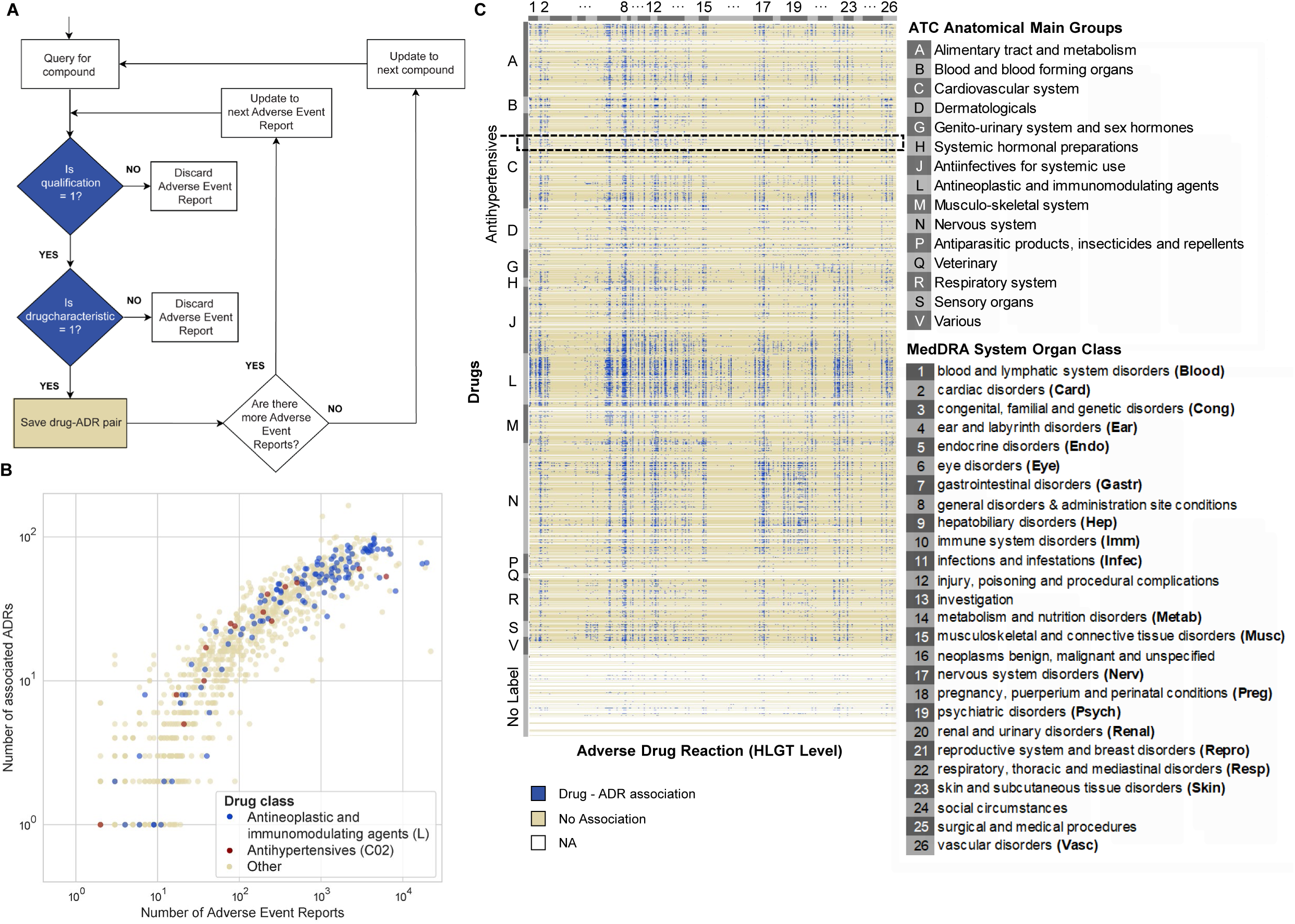
Retrieval of Adverse Event Reports from the FDA Adverse Event Reporting System (FAERS) database. A. **Flow chart of the programmatic strategy for Adverse Event Report retrieval from FAERS by using openFDA.** ‘is qualification = 1’ is a positive filter for adverse event reports that were reported by physicians. ‘is drugcharacterization’ = 1 is a positive filter for drugs that are annotated as the primary suspect drug, which hold a primary role in the cause of the adverse event. B. **Scatter plot of the number of associated ADRs for drugs as a function of the number of adverse event reports retrieved for each drug (N**_**drugs**_ **= 1329).** Drugs without any reported ADR are not shown. C. **Heatmap of ADR profiles (discretized as used for input of random forest model) for all marketed drugs used in this study (N**_**drugs**_ **= 2134).** Drugs are clustered (vertically) according to their ATC drug classes (A-V, or No label if without any ATC code) and HLGT (high level group term) ADRs are grouped (horizontally) according to the parent System Organ Class (SOC) level listed in the legend.

### Random forest model learns relationship between *in vitro* pharmacology and reported ADRs in humans

We deployed a machine learning approach to predict ADRs for a given drug from their *in vitro* secondary pharmacology profiles (Figure 3A). We consider this a multi-label classification problem because a given drug can cause multiple ADRs based on its possible engagement with multiple targets and because a single target may be associated with multiple ADRs. We discretized and one-hot encoded our “input” *in vitro* pharmacology assay data (AC_50_ values) into 3 classes: highly active (AC_50_ < 3 μM), active (3 μM ≤ AC_50_ ≤ 30 μM) and inactive (AC_50_ > 30 μM), which reflect commonly used ranges in the field ^4^. In total, 413 features (assay information) were used to predict 321 High Level Group Term (HLGT) ADRs or 26 System Organ Class (SOC) ADRs for each drug. The observed drug-ADR associations from FAERS, as described above, constitute the dependent variable that the model is learning. We constructed a unifying binary relevance random forest model, which consists of 321 random forest HLGT ADR models. The models were first trained and tested, using 5-fold cross validation where each fold is selected sequentially (Figure 3B). We used 1329 drugs for model construction because these drugs had at least 1 adverse event report in FAERS Q4_2018. The remaining 805 drugs, which did not have any ADR reports, were excluded from training and cross-validation. We confirmed that the distribution of drug classes in our training set (1329 drugs) is comparable to the distribution of drug classes in each 5-fold cross validation split (1063 drugs for training and 226 drugs for testing; Supplementary Figure 1A, χ^2^-test: p-value > 0.99). The model predictions are in probability format, which is used later for target-ADR predictions, and in boolean format (Figure 3A), to enable assessment of model performance via accuracy; macro-precision; macro-recall; Matthew’s correlation coefficient (MCC), a performance measure that takes class imbalance into account; and the area under the receiver operating characteristic curve (macro-AUROC) (Figure 3B). The unifying random forest model performance of SOC ADRs and HLGT ADRs using the full training set (1329 drugs) and the 5-fold cross validation sets (266 drugs, averaged) are depicted in Figure 3B. Accuracy ranges from 0.82 to 0.98, macro-precision ranges from 0.5 to 0.85, macro-recall ranges from 0.29 to 0.74, MCC ranges from 0.37 to 0.83, and macro-AUROC ranges from 0.80 to 0.96. Compared to SOC level (21 ADR terms), the finer grain HLGT level (321 ADR terms) had proportionally fewer drug-ADR associations; additionally, the performance of the HLGT and SOC models are comparable. We therefore proceeded with the HLGT level models for further investigation.

**Figure 3.**
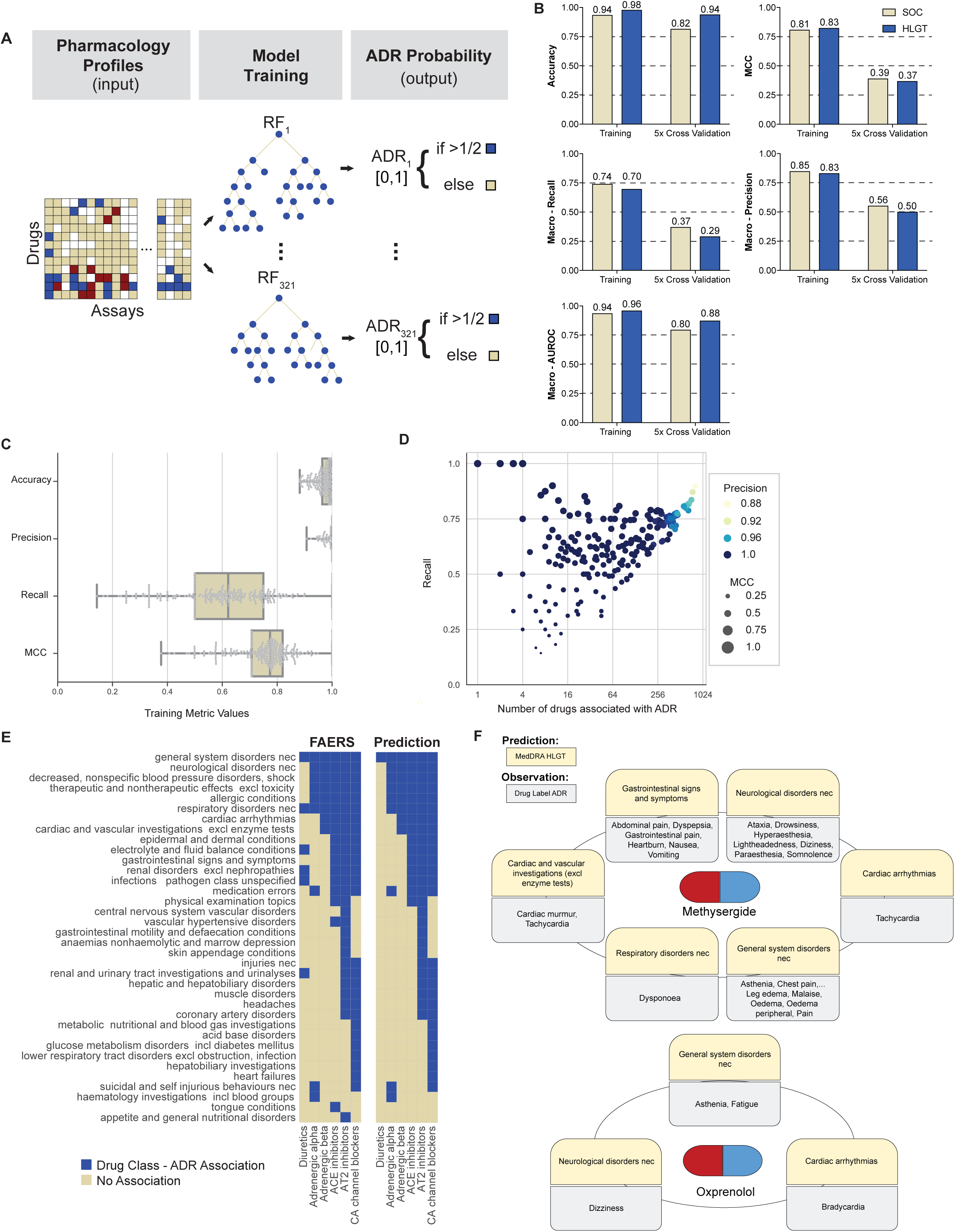
Application of the random forest model to characterize drug-ADR associations. A. **Schematic representation of the machine learning approach.** Using input data, which is a discretized AC_50_ *in vitro* pharmacological profile, we built a separate random forest model for each adverse drug reaction (ADR) that predicts the probability of a drug causing that ADR. For training we used all drugs for which we could retrieve FAERS Q4_2018 adverse event reports (N_drugs_ = 1329). B. **Summary statistics of overall model performance.** We developed two unified random forest models based on two hierarchical levels of organ class specifications. The high level group term (HLGT; blue) unified random forest model consists of 321 ADR random forest models whereas the system organ class (SOC; yellow) unified random forest model consists of 26 ADR random forest models. The performance of the HLGT and SOC models is similar, except in few cases when the HLGT model outperforms the SOC model. (MCC: Matthew’s correlation coefficient, AUROC: area under receiver operating characteristic). Training reflects performance after model training on all 1329 drugs (see A). 5-fold cross validation results are averaged over each fold (all metrics for each fold are detailed in Supplementary Table 4). C. **Box plots indicating the distributions of the training performance metrics** (as in B) for all random forest models of each individual HLGT ADR (N_ADRs_ = 266; center line, median; boxlimits, 1^st^ and 3^rd^ quartiles; whiskers, minimal and maximal value; points represent all data). D. **Scatter plot of the random forest models’ recall (all metrics as in C) as a function of number of associated ADRs**, which served as positive training examples. Colors indicate model precision and circle size reflects the MCC. E. **ADR predictions for anti-hypertensive drugs with different pharmacological targets**. For a set of 22 antihypertensive drugs, we visualized the association between the drugs and HLGT-level ADRs (left). Using the ADR random forest models, we predicted the differences in ADR associations between antihypertensive drugs representing various pharmacological targets (right; overall 36 of the HLGT terms are visualized). True negative predictions (285 HLGT-level ADRs) were omitted from this visualization. F. **Examples of model validation using methysergide and oxprenolol.** The random forest model predicted associations of methysergide with 6 of 321 HLGTs (yellow) which were validated by comparison of ADRs from its drug label (grey) using the SIDER database. One or more of the ADRs corresponding to each HLGT category were confirmed in the drug label.

For 55 of the 321 HLGT ADRs, the corresponding random forest models simply predicted zero for all drugs as mostly none (and at most 4) of the 1329 drugs with adverse event reports were associated with those ADRs (Supplementary Table 3). Since these models were not predictive, we did not consider them for further analyses. For the remaining 266 ADRs, we could determine performance metrics (Figure 3C). Accuracy and precision were high, ranging between 0.9 and 1, whilst the recall and MCC range more widely (Figure 3C). This variability occurs for ADRs that have only a few drugs associated with them (Figure 3D). As the number of associated drugs increases, the models learn to better distinguish true positives from false negatives, subsequently leading to an increase in recall and MCC values (Figure 3D).

### Predictive power of the random forest model for multiple FAERS reporting time periods

To test if our random forest model framework is sensitively dependent on the FAERS reporting period, we constructed new random forest models and performed 5-fold cross validations for both SOC and HLGT levels using FAERS data from 2 different time points: Q4_2014 (including all reports from January 2004 to December 2014) and Q2_2019 (including all reports from January 2004 to June 2019). For proper comparison, the model constructions and cross validations were identical to our above described “main” model based on FAERS Q4_2018. Overall, the performance metrics (accuracy, MCC, macro-precision, macro-recall, macro-AUROC) of both SOC and HLGT level models are comparable between Q4_2014, Q4_2018 and Q2_2019 (Supplementary Table 4). This analysis demonstrates that our random forest modeling framework has a comparable predictive power despite changes in the FAERS reporting time period; therefore, it is not sensitive to different versions of FAERS.

### Chronological validation of predicted drug-ADR associations

To validate the predictive power of our random forest modeling framework further, we performed a chronological validation analysis, through identification of initial false predictions (false positives and false negatives) from the random forest model trained on FAERS Q4_2014 which was then validated using a dataset from the subsequent time period, 2015-2019. The random forest model trained on Q4_2014 data has 421 (0.1% of a total of N=433,671 model predictions) false positive drug - ADR associations, i.e. based on a drug’s pharmacology profile, the model predicted a probability > 0.5 (Figure 3A) for an ADR even though there was no association observed from the adverse event reports up until 2014. However, when compared to the observed Q2_2019 FAERS data, which also include adverse event reports from the time period 2014-2019, 3.1% (13) of the false positives turned into observed drug-ADR associations (true positive), which is 4.4-fold more than expected by chance (χ^2^-test: p-value = 2×10^−5^). Similarly, the Q4_2014 random forest model made 8519 false negative predictions, of which 2.2% (184), 40-fold more than expected by chance (χ^2^-test: p-value < 10^−16^), turned into true negative predictions when compared to the Q2_2019 observed drug-ADR associations. This analysis indicates that significant proportions of our model predictions on drug-ADR associations that were initially “false predictions” became “true predictions” through accumulation of new adverse events reports over time.

### Random forest model predicts expected ADR profiles for anti-hypertensive drugs

As another demonstration of model validation, we analyzed the ADR profiles of 6 subclasses of antihypertensive drugs: adrenergic alpha, adrenergic beta, ACE inhibitors, angiotensin AT2 inhibitors, calcium channel blockers and diuretics (Supplementary Table 5). The signature of the anti-hypertensive drug subclass represents a set of ADRs that were common to all drugs in this subclass. Each antihypertensive drug subclass has a unique ADR fingerprint in the Q4_2018 FAERS version which was closely predicted by our random forest model (Figure 3E). The accuracy ranged from 0.984 to 1, with perfect specificity and precision (Supplementary Table 6). The sensitivity ranged from 0.882 to 1, except for the diuretics sub-class, which had a sensitivity of 0.167. This may be because diuretics target the kidney, and not the cardiovascular system as the rest of the anti-hypertensive drugs do. Of note, the adrenergic alpha and adrenergic beta receptor subclasses maintain distinct profiles in the predicted data. Specifically, the model correctly predicts that adrenergic alpha receptor drugs are associated with suicidal and self injurious behaviors, which has been reported in the literature ^26,27^.

### Random forest model validation through comparison with drug label ADRs

To demonstrate the predictive power of our random forest model on a test set of drugs that were not used for model construction, we utilized the model to predict drug-ADR associations for 805 drugs that did not have any reported ADRs in the FAERS Q4_2018 version, either because there was no match with the drug name or there were no ADR reports for that drug submitted to FAERS by October 2018. For validation, we queried the Side Effect Resource (SIDER) database ^28^, which contains drug-ADR pairs extracted from FDA drug labels by text mining ^28^. For these 805 drugs, we obtained 95 drug matches, which were further reduced to 75 drugs that did not share active ingredients with drugs in the training set. Overall, 57% of positive drug-ADR pairs (i.e. drugs where the model predicts ADRs) were reported in SIDER, compared to 9% of negative pairs (N = 24075; χ^2^-test: p-value < 10^−16^; Supplementary Table 7). For instance, methysergide, a 5-HT receptor antagonist used to treat migraine and cluster headaches, has predicted ADRs from 6 HLGT categories, all of which are supported by specific ADRs from SIDER (Figure 3F). “Cardiovascular disorders with murmurs” appears in the Warnings and Precautions section of the label. Other adverse events under gastrointestinal symptoms and CNS symptoms from SIDER were confirmed in the Adverse Events section. Oxprenolol, a lipophilic beta blocker used for treating angina pectoris, abnormal heart rhythms and high blood pressure, has predicted ADRs from 3 HLGT categories. The specific SIDER ADRs of bradycardia, dizziness and asthenia were also confirmed in the label from the Electronic Medicines Compendium (https://www.medicines.org.uk/emc/product/3235; accessed 09/11/2019). Overall, our random forest model proves to be a powerful tool to predict both on- and off-target related drug-ADR associations from *in vitro* pharmacological drug profiles.

### Random forest model predicts 221 target-ADR associations

To predict which target genes are associated with which ADRs, we utilized the Gini importance score to rank features for their importance in random forest models for each ADR (Figure 4A). For a given ADR, we selected assays that had multiple AC_50_ features represented in the top 5% of Gini scores ranking (see Methods for detail). We then generated ADR probability predictions for an *in silico* compound that is assumed to target only the selected assay with an AC_50_ value corresponding to a represented feature. We also assumed no available data for all other assays. Using this *in silico* AC_50_ profile as an input to the ADR model, we could generate the ADR probability. By assessing differences in ADR probabilities (two sample t-test, FDR corrected p-value < 0.1) between different AC_50_ classes, e.g. highly active (0-3 μM) vs inactive (>30 μM), we predict positive or negative correlations, collectively termed associations, between the selected target assay and ADR. Unsurprisingly, some ADRs did not generate any target associations.

**Figure 4.**
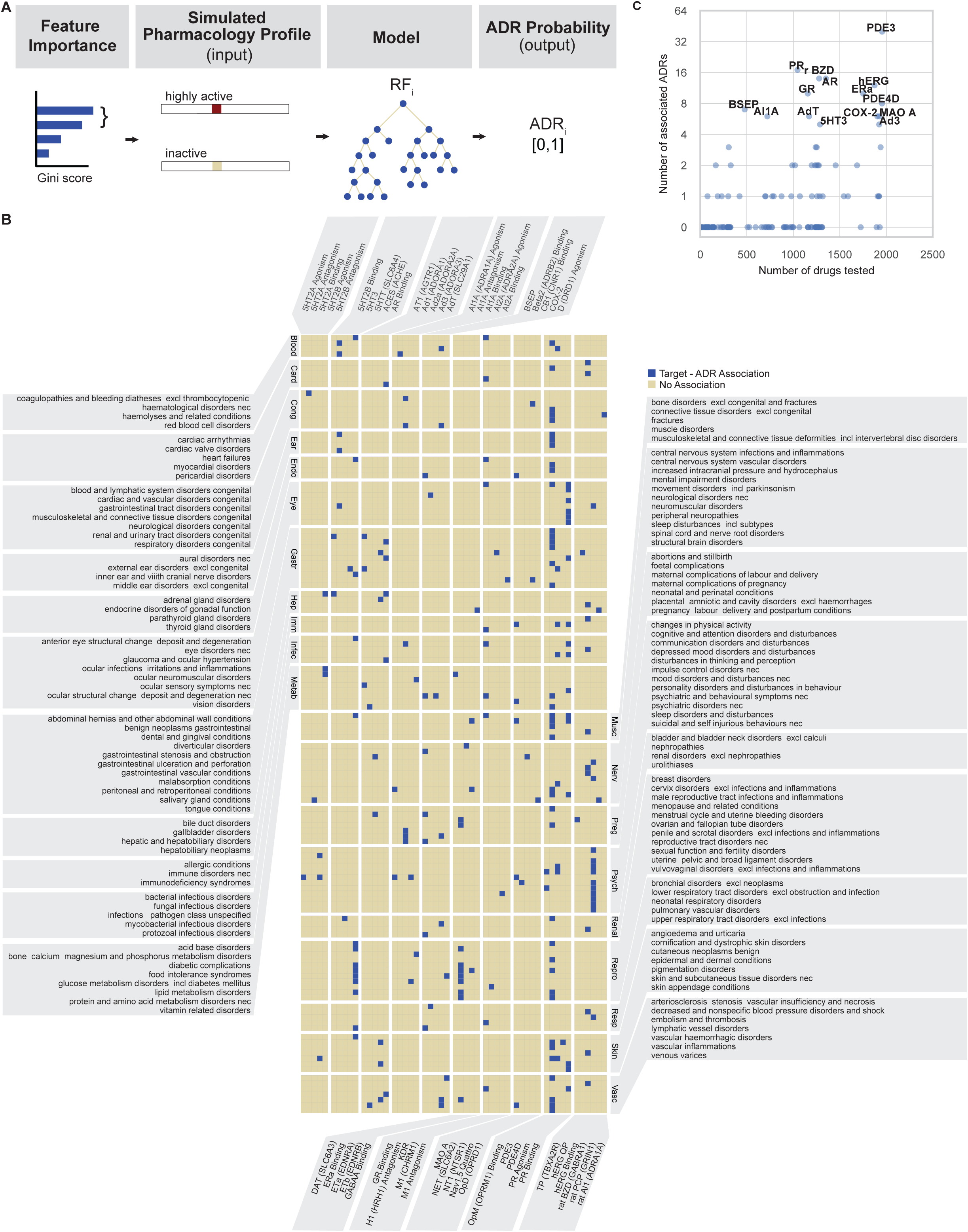
Random forest model predicts target-ADR associations. A. **Schematic outline of the *in silico* ADR-target predictions**. For an ADR of interest, we determined the top 5% of features from the corresponding trained random forest model, ranked according to their Gini importance scores, which measures their contribution to the predictive power of the model. If at least two features (e.g. as depicted: highly active and inactive) from the same target assay are within that top 5%, we determined the ADR probabilities for the simulated cases where an *in silico* compound would target those assay AC_50_ classes only. The ADR probabilities of those simulated cases can then be compared to determine the concentration dependence of the ADR probability. If there is a non-zero correlation between AC_50_ values and ADR probabilities, we conclude that there is an association between the respective ADR and target. For full details, see the Methods. B. **Heatmap showing the resulting 221 predicted target-ADR associations (blue)**. Target (gene symbol) assays are listed alphabetically (horizontal), and HLGT ADRs (vertical) are grouped according to their parent SOC level (as detailed in Figure 2C). For a full description of all target-ADR associations and their ADR probabilities, see Supplementary Table 8. C. **Scatter plot of each target (assay, N=184) showing the number of ADR associations as a function of number of assayed drugs.**

To find biologically meaningful associations, we first filtered out HLGT terms belonging to SOC classes that are not specific to human body parts or only procedural or intervention related (see Methods for detail). Secondly, we filtered out terms that fall under the SOC class neoplasms, since genes are often severely misregulated in cancers and therefore not representative of neoplasm-related ADRs in the organ where the tumor resides. After filtering, we found 221 statistically significant target-ADR associations (Figure 4B, full details including p-values in Supplementary Table 8); 51 out of 184 target assays and 132 out of 321 ADRs are represented (Figure 4B). The assay class distribution of these 51 targets, represented among the 221 predicted target-ADR associations, is similar to the class distribution of all target assays (Supplementary Figure 1B, χ^2^-test: p-value = 0.09). This demonstrates that our algorithm does not bias towards certain target classes. In the following sections, we investigate these 221 target-ADR associations in more detail.

### Systematic literature validation of target-ADR associations

To validate our ADR-target predictions, we performed a systematic literature co-occurrence analysis. First, we mapped all genes corresponding to the assays and HLGT level ADRs to their respective MeSH terms (Supplementary Table 9). Next, we queried PubMed for the publication identifiers linked to these MeSH terms and determined the number of publications that corresponded to both a gene and HLGT term (i.e. co-occurrence). We found at least one co-occurrence publication for 66% (145) of 219 predicted unique gene-HLGT MeSH pairs, which was higher (Fisher Exact test: odds ratio=1.8, p-value=6×10^−5^) than for all possible negative unique gene-HLGT pairs (N=26705). In order to control for the fact that some ADRs and genes are studied more intensively than others, we also compared our set of positive predictions to a negative control set (N=4890) formed by permuted pairs from the positive set and obtained similar results (Fisher Exact test: odds ratio=1.5, p-value=3×10^−3^). Furthermore, as quantified by the co-occurrence “lift” over the reporting probability when assuming independence,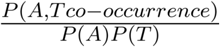 (see Methods for details), we found 4-fold higher co-occurrence median lift values for our predictions compared to all negative pairs (Mann Whitney U-test: p-value=2×10^−5^), and 3-fold higher lift than permuted negative pairs (Mann Whitney U-test: p-value=3×10^−4^). We conclude that our target-ADR identification method provides association predictions that are supported by the literature in higher proportion than random selection of target-ADR pairs.

### Evidence for targets that are predicted to cause cardiovascular-related ADRs

To further validate our model’s ability to predict target-ADR associations, we investigated a group of cardiovascular ADRs. We found that hERG binding was associated with cardiac arrhythmias and heart failure (Table 1). hERG encodes for a subunit of the cardiac potassium ion channel and contributes to cardiac electrical activity, which is necessary to regulate the heartbeat. The mechanism of action for drug-induced arrhythmias by blocking hERG has been described in numerous human ^29^ and animal studies ^30^, as well as structural modeling ^31^ studies (Table 1). Consistently, our systematic PubMed queries found 753 co-occurrence publications in support of this predicted association and 6 co-occurrences for hERG binding increasing the risk of heart failure. We did not find an ADR probability associated within the range of 0-3 μM AC_50_ of hERG binding, likely because such strong binding to hERG is a common reason for deprioritizing drug candidates in development ^32^.

**Table 1:** Predicted associations between targets and cardiac ADRs. High Level Group Terms (HLGT; MedDRA) associations with targets and Adverse Drug Reaction (ADR) probability in three concentration ranges (third column). Evidence of the ADR-target pairs were obtained from peer reviewed publications (fourth column). The number of publications linked to both an HLGT ADR and target gene was obtained via a systematic literature co-occurrence analysis (fifth column). hERG: human Ether-a-go-go-Related Gene associated potassium channel; PDE3: phosphodiesterase-3 enzyme; GR: glucocorticoid receptor; AdT: Adenosine transporter; COX-2: cyclooxygenase enzyme, type 2.

| Cardiac Disorder HLGT | Target | ADR Probability |  |  | Literature evidence<br>human (h), animal (a), *in vitro* (v) | Co-occurrence<br>(number) |
| --- | --- | --- | --- | --- | --- | --- |
|  |  | 0-3 µM | 3-30 µM | >30 µM |  |  |
| cardiac arrhythmias | hERG (Binding) | - | 0.03 | 0.002 | h<sup>29</sup> a<sup>30</sup> v<sup>31</sup> | 753 |
| cardiac valve disorders | PDE3 | 0.05 | - | 0 | h<sup>33,34</sup> | 3 |
| heart failures | hERG (Binding) | - | 0.005 | 0 | h<sup>63</sup> | 6 |
| myocardial disorders | GR (Binding) | 0.02 | - | 0.005 | h<sup>36,37</sup> | 8 |
| pericardial disorders | AdT | - | 0.01 | 0 | a<sup>35</sup> | 0 |

The model predictions also suggest that PDE3 inhibition is associated with cardiac valve disorders (Table 1, 3 co-occurrence publications). PDE3 inhibition is used clinically to treat dilated cardiomyopathy ^33^, which encapsulates valvular heart disorder. However, the PDE3 therapeutic window is narrow, partially due to complex signaling networks ^34^, and careful dosing is required to avoid increased mortality in response to treatment.

**Table 2:** Predicted renal ADR - target associations (detailed legend in Table 1).

| Renal Disorder HLGT | Target | ADR Probability |  |  | Literature evidence<br>human (h); animal (a), *in vitro* (v) | Co-occurrence<br>(number) |
| --- | --- | --- | --- | --- | --- | --- |
|  |  | 0-3 µM | 3-30 µM | >30 µM |  |  |
| nephropathies | COX-2 | 0.003 | - | 0 | h<sup>38</sup> a<sup>39,40</sup> | 398 |
| renal and urinary tract disorders congenital | PDE3 | 0.004 | - | 0 | h<sup>56,64</sup> a<sup>41,65</sup> | 0 |
| renal disorders excl nephropathies | hERG (Binding) | - | 0.01 | 0.0007 | h<sup>42</sup> a<sup>66</sup> | 2 |

**Table 3:** Predicted ADR associations with inhibition of the Bile Salt Export Pump (BSEP) transporter (detailed legend in Table 1).

| HLGT | Target | ADR Probability |  |  | Literature evidence | Co-occurrence<br>(number) |
| --- | --- | --- | --- | --- | --- | --- |
| | | 0-3 $\mu\text{M}$ | 3-30 $\mu\text{M}$ | >30 $\mu\text{M}$ | human (h); animal (a) | |
| central nervous system vascular disorders | BSEP | - | 0.09 | 0.008 | for BSEP and bile acid) a<sup>67</sup> | 2 |
| foetal complications | BSEP | 0.01 | - | 0 | h<sup>50</sup> | 7 |
| pregnancy labour delivery and postpartum conditions | BSEP | - | 0.1 | 0 | h<sup>51</sup> | 0 |
| lipid metabolism disorders | BSEP | - | 0.2 | 0 | h<sup>68,69</sup> | 5 |
| thyroid gland disorders | BSEP | - | 0.07 | 0 | a<sup>70,71</sup> | 1 |
| upper respiratory tract disorders excl infections | BSEP | 0.1 | - | 0 | h<sup>72</sup> a<sup>73</sup> | 0 |
| urolithiasis | BSEP | - | 0.07 | 0 | h<sup>74</sup> | 0 |
| | | 0-30 $\mu\text{M}$ | 30-300 $\mu\text{M}$ | >300 $\mu\text{M}$ | | |
| hepatic and hepatobiliary disorders | BSEP | - | 0.2 | 0.09 | h<sup>48</sup> | 354 |

Furthermore, our model predicts that adenosine transporter (AdT) inhibition increases the risk of pericardial disorders (Table 1). For this scenario, we did not find direct supporting evidence in the literature, however there is evidence that disturbed adenosine homeostasis in pathological cardiac conditions could result in pericardial effusion or pericarditis ^35^.

The model suggests that glucocorticoid receptor (GR) binding is more likely to lead to myocardial disorders if the drug has high affinity for GR (Table 1, 8 co-occurrence publications). This is supported by the finding that glucocorticoid treatment of patients with rheumatoid arthritis increased the risk of myocardial infarction ^36^. Furthermore, it is known that dysregulation of glucocorticoids can give rise to cardiotoxicity ^37^.

Taken together, this investigation of genes associated with cardiovascular ADRs confirms the well-known association of hERG with cardiac arrhythmia, and also highlights ADR associations that would merit further experimental investigation.

### COX-2, PDE3, and hERG associations with kidney related ADRs

Another important class of ADRs involve the kidney (Figure 4B, label: renal). We found COX-2 associated with nephropathies (Table 2), which has been well recognized (398 co-occurrence publications) and evidenced previously ^38–40^. Interestingly, another model prediction is PDE3 sensitivity correlating with congenital renal and urinary tract disorders (Table 2). According to a mouse model study ^41^, PDE3 inhibition could be a contributing factor in Polycystic Kidney Disease (PKD), as PDE3 protein levels are downregulated in PKD compared to healthy control kidneys. Lastly, we found an unexpected association between hERG and renal disorders (excluding nephropathy) (Table 2). One study has found a loss of hERG function in renal cell carcinoma ^42^. In humans, hERG expression in the kidney is much lower than in the heart ^43^. Therefore, we conclude that a link between hERG and renal disorders remains a prediction that warrants further investigation.

### PDE3 and nuclear hormone receptors AR, ERa, and PR are overrepresented in ADR associations

To investigate if the number of different drugs tested for a target assay is predictive to the number of ADRs associated with that target (Figure 4C), we calculated their Spearman correlation coefficient and found a moderate correlation (ρ=0.5; Figure 4C). However, some targets had considerably more associated ADRs than other targets that were tested a similar number of times, indicating that more frequently performed assays do not necessarily result in a higher number of associated ADRs (Figure 4C). Out of all target assays, PDE3 was associated with the most ADRs (Figure 4C), falling in a wide range of SOC classes (Figure 4B, Supplementary Table 8). Furthermore, nuclear hormone receptors for androgen (AR), estrogen (ERa) and progesterone (PR) binding assays also have disproportionately many ADR associations, compared to their frequency of testing (Figure 4C). As expected, AR (7/14 ADRs), ERa (9/10 ADRs) but not PR (0/17 ADRs) are associated with sexual reproductive organ- and pregnancy-related ADRs (Figure 4B, Supplementary Table 8). Androgen is produced in the adrenal gland ^44^ and we predict a link between AR with adrenal gland disorders, with evidence in mouse studies ^45^. Interestingly, the model predicted 6 ocular ADRs associated to PR, including vision disorders, anterior eye structural change (deposit and degeneration), infections, irritations and inflammations and structural changes (Figure 4B, Supplementary Table 8), for which we could find supporting evidence ^46^.

### GABA_A_ receptor associations with psychoactive ADRs

GABA_A_ receptor is the primary target of benzodiazepines (BZD), a drug class known to be psychoactive with potential of addiction ^47^. Consistently, our model predicts that this ligand-gated chloride ion channel assay is associated with 14 ADRs, 13 of which are neurologically and psychiatrically related, including disturbances in thinking and perception, sleep disorders, depression and suicidal behaviors (Figure 4B, Supplementary Table 8).

### Bile salt export pump BSEP associations with ADRs in various organs

BSEP, encoded by *ABCB11* and a member of the superfamily of ATP-binding cassette (ABC) transporters, is most highly expressed in the liver ^43^. Drugs that target BSEP are often associated with hepatotoxicity ^48^. However, initially, we did not find a BSEP association with hepatic and hepatobiliary disorders. To investigate this false negative prediction, we noted that the dynamic range of the BSEP assay specifically extends up to 300 μM because the first pass effect for orally delivered drugs results in high concentrations in the liver ^49^; as a result, most of our data falls into the ‘inactive’ (>30 uM class). Consistently, the BSEP inactive feature had the highest Gini score for this HLGT term, while its two active features had much lower Gini scores, falling outside of the top 5%. To take the extended dynamic range into account, we altered the BSEP assay class boundaries to 0-30 μM, 30-300 μM and >300 μM and retrained the random forest model. In this case, we did find BSEP associated with hepatic and hepatobiliary disorders (Table 3, 354 publication co-occurrences), according to our association criteria (Figure 4A). We repeated this procedure whilst replacing the first class boundary (30 μM) with 100 μM and found the same association again, indicating the robustness of our results. Interestingly, with our original AC_50_ discretizations (Figure 1D), we found BSEP associated with 7 other ADRs from various organ classes (Table 3), much more than other targets that were assayed at a similar frequency (Figure 4C). This suggests that compounds potent against BSEP (AC_50_ < 30 μM) could cause adverse effects in addition to hepatotoxicity, which already occurs at lower potency. We found BSEP associated with urolithiasis and with disorders of the thyroid gland, upper respiratory tract disorders (excl infections), lipid metabolism and central nervous system (Table 3). Since BSEP expression is much lower in these organs ^43^, we searched the literature for evidence including a substrate of BSEP, bile acid. We could find previous studies linking bile acid to these disorders (Table 3), which suggests an indirect relation between BSEP and these ADRs through bile acid metabolism. Lastly, we found BSEP associated with foetal complications and pregnancy conditions (Table 3), both supported through prior studies that link BSEP with transient neonatal cholestasis and intrahepatic cholestasis of pregnancy, respectively ^50,51^.

## Discussion

In this study we have taken a machine learning approach to predict human ADRs from the *in vitro* secondary pharmacology profiles of a large number of marketed and withdrawn drugs. Several prior studies focus on predicting ADRs directly from chemical drug structure ^52,53^. However, utilizing functional information such as *in vitro* pharmacological targeting of common (off) targets represents a viable alternative to bridge the complex relationship between drugs and their effects in the human body ^4^.

Our random forest model performance metrics are good considering the sparse coverage (2134 drugs) over a large input space (3^184^ possibilities) and partial overlap with ADR reporting for these drugs, making ADR occurrence prediction effectively a one shot learning task. Importantly, our model performances were strong enough to discover drug-ADR and biologically meaningful target-ADR associations. To determine the target-ADR associations, we utilized all available input data for model training and made use of Gini scores to robustly select relevant features for ADR probability predictions. We rigorously validated our model predictions with multiple independent analyses (e.g. chronological validation on drug-ADR associations and systematic literature validation on target-ADR associations). Our novel method for target-ADR associations was able to recapitulate well recognized causal relations, such as hERG with cardiac arrhythmias. For others, we were able to find literature evidence in animal or *in vitro* studies but our study is, to our knowledge, a first in human report. Another fraction of target-ADR associations represents predictions of novel, unexpected or little known associations, such as Adenosine Transporter (AdT) and pericardial disorders, for which we could find little evidence other than our analysis of adverse event reports. Similar to genome-wide association studies, our quantitative methodology extracts statistically significant relations from human population data. With this framework in mind, our 221 associations form a rich resource that can be used for further mechanistic studies in the drug discovery process.

Our random forest model is agnostic to molecular mechanisms; therefore, resulting associations could arise from indirect regulation. A likely example is the bile transporter BSEP, which is associated with numerous ADRs, although it is most highly expressed in the liver and kidney. We have related our findings to evidence that misregulation of its substrate, bile acid, could result in disorders related to kidney stones, lipid metabolism, thyroid gland, respiratory system, and central nervous system. This also indicates the strength of our approach, which can relate genes to physiological processes unbiasedly in humans, without any interventions or large scale genome-wide association studies, but solely with voluntary adverse event reporting.

Some of the predicted target-ADR associations could be hard to validate, such as the PDE3 enzyme association with congenital renal disorders association. While the association is valid, the modality has to be clarified: PDE3 inhibitors are proposed to ameliorate certain forms of chronic kidney disease ^56^, instead of causing it. Thus, predictions of congenital disorders should be considered but confirmed by checking the modality of the effects.

While we recommend this approach to find target-ADR associations to impact safety awareness in drug discovery, we are also aware of the limitations. Firstly, targets in the *in vitro* pharmacology panel cover a fraction of the biological target space and not all drugs were tested in all assays. We recognize that 47% of all targets belong to the GPCR target family with limited representation of other therapeutic or ADR-associated targets such as ion channels and kinases. However, our model predictions are not biased towards the GPCR target family; the target classes of 51 targets in 221 target-ADR association predictions have a similar distribution compared to all target classes in our input data (Supplementary Figure 1B). Also, data are influenced by prior knowledge; for example, more than 87% of all drugs in the set were tested for hERG activity. High affinity (lower AC_50_ value) for hERG is associated with higher probability for QT prolongation for human and non-human preclinical species ^29,30^. As discussed earlier, there are not many drugs with a hERG AC_50_ value in the highly active class (0-3 μM), which is a commonly encountered roadblock for drug candidates to progress towards clinical trials ^32^. Only about 10% of all drugs fall into the highly active class in our assay data. To limit feature engineering, our AC_50_ discretization into three classes (Figure 1D) was kept uniform across all assays. Notably for the BSEP assay only, the dynamic range extends up to 300 μM and as a result most of our data falls into the ‘inactive’ (>30 μM) class. Consequently, we initially did not find the expected association with hepatotoxicity. We rectified this by reclassifying the BSEP assay data according to levels required for hepatotoxicity of BSEP inhibition ^54,55^ and indeed recovered the expected association.

Secondly, *in vitro* potency is an initial marker of clinical effect, and does not take into account prolonged dosing, comorbidity or pharmacokinetic/pharmacodynamic (PK/PD) relationships (e.g. therapeutic window). For 9 of 184 assays, non-human proteins were assayed (e.g. rat brain was used as a source for the benzodiazepine receptor) which may not be a direct correlate of the human protein. One way of further improvement of our approach is to include additional occupancy parameters and PK/PD components for higher precision and enhanced predictive value.

Lastly, in the FAERS database, drug-ADR associations may be mislabeled, e.g. anti-hypertensives are often reported as associated with hypertension as an ADR, rather than as the indication. Additionally, the FAERS database does not contain information on the total number of patients exposed to a particular drug, nor is it necessarily a reflection of the true incidence or frequency of ADRs. These and other limitations are discussed by Maciejewski et al. ^21^ with suggestions and methodology for further refinement of the FAERS database curation and maintenance.

We investigated one-to-one associations between targets and ADRs because these relationships are biologically meaningful and have a high utility in preclinical drug development. However, in some cases, a given ADR can be a prerequisite for others (e.g. hypotension leading to reflex tachycardia). We leave a model extension to incorporate these dependencies as future work. For target-ADR associations, we utilized our random forest model for a single drug at a time. Our model can be repurposed to predict possible ADRs from combination drug therapies and likelihood of drug-drug interactions. In principle, this can be extended for combination therapies by merging the *in vitro* data from the individual compounds and the predictions can be validated by querying Offside and Twosides databases ^57^. Similarly, our model can be utilized for drug repositioning and repurposing, using similar drug-ADR and target-ADR profiles. In conclusion, our random forest model and the target-ADR associations provide a validated, comprehensive resource to support drug development and future human biology studies.

## Methods

### *In vitro* secondary pharmacology assays for marketed drugs

AC_50_ values of 2134 marketed drugs (Supplementary Table 1) were measured in up to 218 different *in vitro* secondary pharmacology assays. Compounds were obtained from the Novartis Institutes of Biomedical Research (NIBR) compound library and tested in a panel of *in vitro* biochemical and cell-based assays at Eurofins and at NIBR in concentration-response (8 concentrations, half-log dilutions starting at 30 µM). Assay formats varied from radioligand binding to isolated protein to cellular assays. Example protocols may be found at https://www.eurofinsdiscoveryservices.com/cms/cms-content/services/in-vitro-assays/. Normalized concentration response curves were fitted using a four parameter logistic equation with internally developed software (Helios). The equation used is for a one site sigmoidal dose response curve Y as a function of tested concentrations X: Y(X)=A+(B-A)/(1+(X/C)^D^), with fitted parameters A=min(Y), B=max(Y), C=AC_50_ and exponent D. By default, A is fixed at 0, whereas B is not fixed.

If a drug was not tested against a specific assay, the AC_50_ value was set to NA (not available). AC_50_ values from similar assays with the same gene target were merged to reduce the NA data and features in the random forest model; this procedure resulted in 184 different target assays (Supplementary Table 2). In case any merged assays had multiple AC_50_ values for the same drug, we averaged these geometrically to take into account variation over orders of magnitudes. In figures 1D and 2C, the drugs are classified according to their annotated Anatomical Therapeutic Chemical (ATC) code ^19^. In case of multiple ATC codes, we assigned the most frequent level 1 code.

### Mining adverse event reports of marketed drugs using OpenFDA

In this study, we utilized openFDA to acquire FAERS reports related to the query compounds ^15,20^. This Elasticsearch-based API provides a raw download access to a large volume of structured datasets, including adverse events reports from FAERS.

We used generic compound names (e.g. “Amoxicillin”) to query through the openFDA interface, accessed programmatically using Python. In order to maximize the coverage over FDA datasets, we normalized generic names to uppercase format followed by a name similarity metric to filter out unrelated records in our analysis. We included reports when the Jaro similarity between the query generic name and reported compound name was equal or greater than 0.8. To illustrate, to query “3alpha-Androstanediol”, we acquired reports including “3α-Androstanediol”, “Androstanediol”, “3-alpha-Androstanediol” as different lexical variations of the generic name and collated the resulting adverse event reports.

As the FAERS database contains information voluntarily submitted by healthcare professionals, consumers, lawyers and manufacturers, adverse event reports may be duplicated by multiple parties per event, and may be more likely to contain incorrect information if submitted by a non-medical professional. To reduce reporting bias and increase report information accuracy, we only analyzed reports submitted by physicians (data field: ‘qualification’ = 1). In this subset of adverse event reports, the data were further filtered by reported drug characterization, which indicates how the physician characterized the role of the drug in the patient’s adverse event. A drug can be characterized as a primary suspect drug, holding a primary role in the cause of the adverse event (data field: ‘drugcharacterization’ = 1); a concomitant drug (‘drugcharacterization’ = 2); or an interacting drug (‘drugcharacterization’ = 3). Here, we included only primary suspect drug reports. Without this restriction, model performances did not improve. We obtained all adverse events reports corresponding to the query compound that passed through the aforementioned filters.

Adverse event report descriptions are coded as medical terms of MedDRA terminology ^17^. Medical observations can be reported using 5 hierarchical levels of medical terminology, ranging from a very general System Organ Class term (e.g. gastrointestinal disorders) to a very specific Lowest Level Term (e.g. feeling queasy). Each term is linked to only one term on a higher level. For each report, we recorded all MedDRA Reaction terms (data field: “reactionmeddrapt”) at the Preferred Term level and mapped these Preferred Terms to Higher Level Group Term and System Organ Class level. For each (ADR term, drug) tuple, we then calculated the ADR occurrence, defined as the following fraction: number of adverse event reports containing that ADR term relative to the total number of ADR reports for that drug.

For different FAERS versions (Q4_2014, Q4_2018 and Q2_2019), we used the same query except the time parameter TO, which was set to 12/30/2014 for the Q4_2014 query. For other two queries, we didn’t set the limit parameter which was filled with the query time by default (query date was 10/10/2018 for Q4_2018 and 08/12/2019 for Q2_2019).

### Random forest models

To construct and train our models (Figure 3A), we used AC_50_ values for a panel of target assays for marketed drugs (model input; independent variable) and ADR occurrences of the compounds (model output/predictions; dependent variable). Since there may be several ADRs associated with any given drug, we considered this a multi-label learning problem. We took a “first-order strategy”, i.e. we assume there is no correlation between different ADRs, and a “divide and conquer” strategy, i.e. we decompose our multi-label learning task into n independent binary classification problems, where n is the number of different ADR terms in our output data (n = 26 for SOC and n = 321 for HLGT level respectively). We built a random forest^58^ binary classifier for each ADR using Binary Relevance with the random forest modeling option in mldr package ^59^ and utiml package in R ^60^.

To define the features for the random forest models, we discretized and one-hot encoded our input AC_50_ values. Discretization was essential to limit the number of features and enhance the predictive power of the model. We defined 3 classes (levels) of AC_50_ ranges for each target assay.

- Highly active class: AC_50_ in [0, 3 μM)
- Active class: AC_50_ in [3 μM, 30 μM]
- Inactive class: AC_50_ greater than 30 μM

If the AC_50_ value is NA, the values for all Classes are 0. Each drug has AC_50_ values for 184 (merged) assays, so there are 184×3 = 552 binary features to represent our input data. Features consisting of only 0 values were removed, resulting in 413 input features used for model construction.

The observed ADR occurrences were discretized into binary dependent variables. To achieve this, first let N_d_ be the total number of ADR reports for a given drug. The probability to observe an ADR occurrence O^ADR^ = X / N_d_ at random is equivalent to choosing that ADR X times out of N_d_ with X distributed binomially: X∼bin(N_d_, p=1/n). Here, n represents the total number of ADRs as defined above. Under this null distribution, we calculate the p-values for all observed ADR occurrences O^ADR^ for a given drug, and then perform a Benjamini-Hochberg False Discovery Rate (FDR) correction (using the Python statsmodels package). If an FDR-corrected p-value is < 0.01, then the ADR value for that drug is 1, reflecting an association; 0 otherwise.

All random forest models were first trained using 5-fold cross validation and each fold is selected sequentially. 1063 drugs were used for training and 266 drugs were used for testing in each fold, and the distribution of drug classes (ATC) in our training set (1329 drugs) is preserved in 5-fold cross validation splits (Supplementary Figure 1A), i.e. the drug classes are represented in 5-fold cross validation splits the way they are represented in our training set. Then, the drugs with at least 1 ADR report are used as a training set. For a given (drug) input of AC_50_ values and ADR, the random forest model output, termed ADR probability, can be interpreted as the probability that the ADR is associated with the drug. To enable direct comparison of model predictions with binarized ADR occurrences, we binarized these ADR probabilities with a simple threshold value of 0.5. These binary values were used for training, cross validation and to calculate classification performance metrics (Figure 3B,C). All models have been constructed the same way regardless of different FAERS versions.

We evaluated our models based on five metrics: accuracy, Matthew’s correlation coefficient (MCC), macro-precision, macro-recall and area under the receiver operating characteristic curve (macro-AUROC). These metrics are calculated using their definitions below, except 2 metrics: (1) MCC, which is calculated using mltools package in R (https://github.com/ben519/mltools) and (2) AUROC, which is calculated using precrec package in R ^61^.

- Accuracy = (*TP* + *TN*) / (*TP* + *TN* + *FP* + *FN*)
- Precision = *TP* / (*TP* + *FP*)
- Recall = *TP* / (*TP* + *FN*)
- MCC = (*TP* * *TN* − *FP* * *FN*) / *SQRT* ((*TP* + *FP*) * (*TP* + *FN*) * (*TN* + *FP*) * (*TN* + *FN*))
- 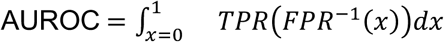

where TPR (true positive rate = *TP* / (*TP* + *FN*)) and FPR (false positive rate = *FP* / (*F P* + *TN*)). The corresponding metrics for each ADR model (Figure 3C, 3D) are accuracy, precision, recall, and MCC, which is calculated using mltools package in R (https://github.com/ben519/mltools).

### Determination of target-ADR associations

To find associations between gene target assays and ADRs (Figure 4), we first generated ADR probabilities specific to a given assay. As a model input, one out of its three random forest input features’ value was set to 1 and all others to 0. This simulates the scenario of an *in silico* compound that is potent with an AC_50_ value in the range corresponding to the positive feature only. We then utilized the ADR’s random forest model, pre-trained on all available marketed drug data (see previous section), to calculate the resulting ADR probability. We repeated this procedure for each feature of all assays and each ADR.

To select the predictive features for a given ADR, we ordered the pre-trained random forest model’s input features according to their Gini importance score ^62^ and denote the top 5% as significant features. Our criteria for a gene (target assay) - ADR pair were:

- For a given ADR: at least 2 out of 3 assay features need to be significant in order to make a reliable comparison between the ADR probabilities with respect to AC_50_ values.
- At least one of the ADR probabilities of the significant features has to be larger than zero.

We filtered out target-ADR pairs if the ADR term maps to the following SOC classes, which are not specific to body parts or underlying human biology:

- general disorders and administration site conditions
- injury, poisoning and procedural complications
- investigations
- neoplasms benign, malignant and unspecified (incl cysts and polyps)
- poisoning and procedural complications
- social circumstances
- surgical and medical procedures

To ensure the reproducibility of the target-ADR pair selection procedure, we repeated the random forest model training with different seeds for a total of 5 times. We then took the union of the 5 sets of target-ADR pairs and discarded pairs that were only found once out of 5 runs. Finally, to determine if the mean ADR probabilities between the selected AC_50_ classes were statistically significantly different, we performed a two-sample t-test with sample sizes equal to the number of times a class was selected (ranging from 2 to 5 times) using the Python scikit.stats package. In case all three AC_50_ classes were represented, we tested the highly active versus inactive class. We then performed a Benjamini-Hochberg FDR correction. If the FDR-corrected p-value is < 0.1, then the target-ADR pair is considered a statistically significant association (Figure 4B, Supplementary Table 8).

To evaluate the relation between the HLGT level ADR term hepatic and hepatobiliary disorders and target assay BSEP, we also trained and analyzed two random forest models as described above to find target-ADR pairs but with only the BSEP assay data discretized with class boundaries [0, 30 μM), [30, 300 μM] and >300 μM or [0, 100 μM), [100, 300 μM] and >300 μM.

### Side Effect Resource (SIDER) analysis

The Side Effect Resource (SIDER; version 4.1) was downloaded (http://sideeffects.embl.de/download/; accessed on 09/16/2019). The file meddra_all_se.tsv.gz contains drug-ADR pairs extracted from drug labels using text mining ^28^. The supplied MedDRA preferred term (PT) was mapped to HLGT used for the random forest modeling. The file drug_atc.txt provides mappings from drug names as used in SIDER to Anatomical Therapeutic Chemical (ATC) codes. ATC codes for the 805 drugs in the test set were obtained from the NIBR compound database, and matched to ATC codes from SIDER. For drugs that could not be matched via ATC codes, additional matches were obtained by mapping the compound name, first trying the name in its entirety (e.g. “butriptyline hydrochloride”, then on the first word in the drug name (e.g. “butriptyline”). All matches, whether obtained on ATC codes or by drug name, were reviewed manually for accuracy.

### Systematic validation of predicted target-ADR association using PubMed database

We built a query based on 254 unique HLGT level ADR terms and 106 unique target genes (corresponding to the assays), for which we could find a corresponding MeSH term (Supplementary Table 9), to retrieve linked publication identifiers (PMIDs) from the PubMed database. All PMIDs were acquired by submitting a query for every MeSH entity separately via the PubMed API engine, a search engine that provides access to the MEDLINE database of references and abstracts on life sciences and biomedical articles. Next, we determined the PMIDs for a gene-ADR pair as the intersection of the two PMID sets of each corresponding MeSH term query. Furthermore, for each possible gene-ADR pair we determined whether it was part of the 221 predicted associations from the Random Forest model or not. In this way, we obtained 219 unique positive gene-ADR pairs and a total 26705 unique negative pairs. Lastly, we generated a set of negative pairs corresponding to all permutation pairs from the 39 unique genes and 131 unique ADRs that are part of the positive set, resulting in 4890 unique negative pairs in this negative control set. To assess any statistical overrepresentation, we calculated the number of pairs with at least one co-occurrence publication for both negative and positive sets and assessed significance with a Fisher Exact test (Python function scipy.stats.fisher_exact). Furthermore, we calculated the co-occurrence “lift” over the reporting probability when assuming independence, defined as 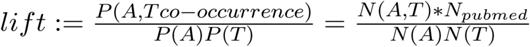, With *N*_*pubmed*_ = 29138919 the total number of PMIDs in the Pubmed database in 2019 (https://www.nlm.nih.gov/bsd/licensee/2019_stats/2019_LO.html). *N*(*A, T*), *N*(*A*), and *N*(*T*) are respectively the number of retrieved PMIDs for a unique gene-ADR pair, ADR, or gene target separately. To assess the location differences of the above described positive versus negative distribution of lift values, we performed a Mann Whitney U test (Python function scipy.stats.mannwhitneyu, two-sided, continuity correction=True).

### Data availability

Data retrieved from public repositories is made available in Supplementary Tables 1-9 and on GitHub (https://github.com/samanfrm/ADRtarget).

### Code availability

Code to query FAERS and PubMed, to construct the random forest models and identify the target-ADR associations is made available on GitHub (https://github.com/samanfrm/ADRtarget).

## Figure Legends

**Supplementary Figure 1.**
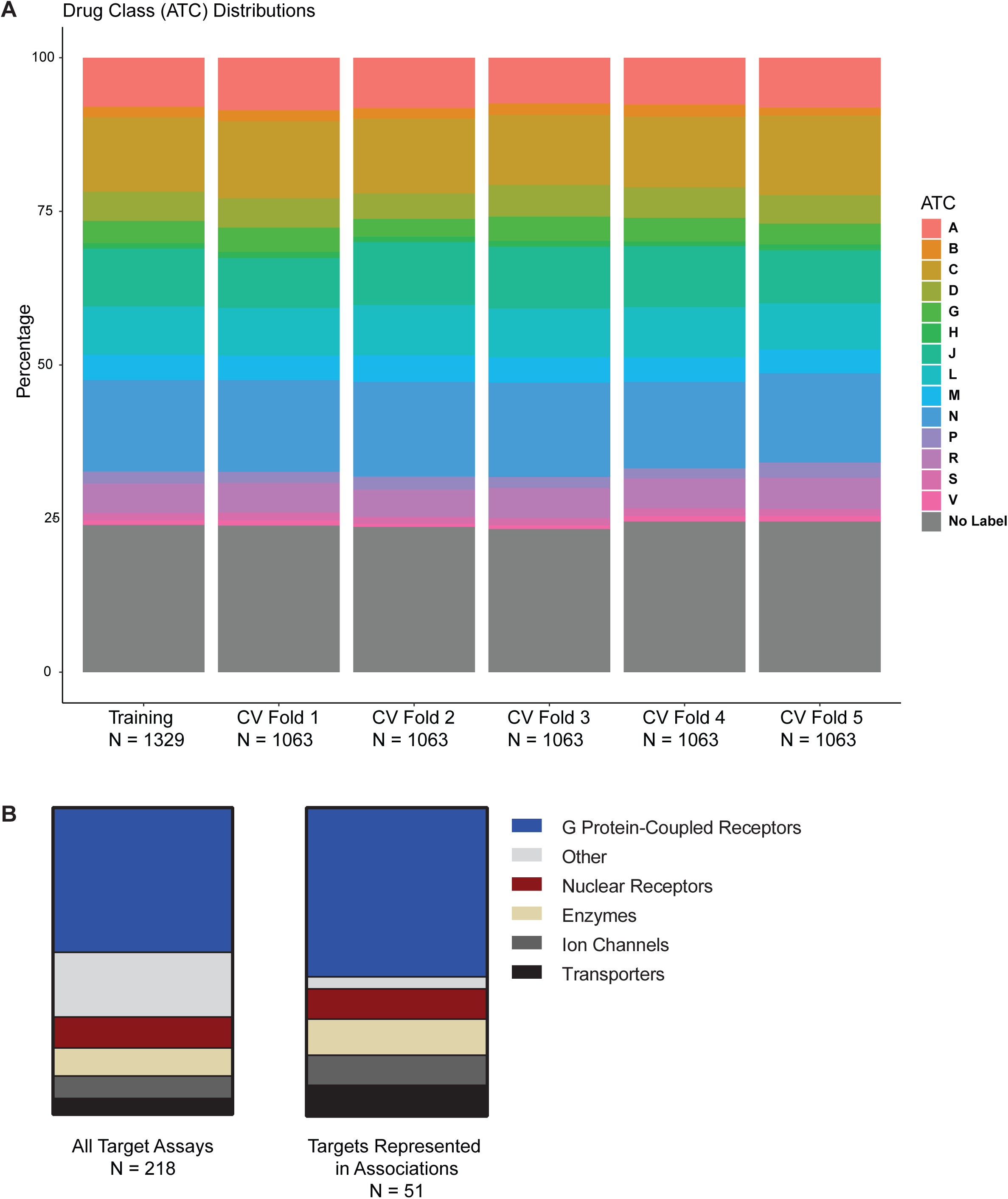
Drug and target class distributions of random forest model. A. **Stacked bar charts comparing drug class distributions of the complete training set (1329 drugs) and each 5-fold cross validation (CV) fold (1063 drugs).** No significant differences were observed: (χ^2^-test: p-value > 0.99). Drug class labels (ATC) are as described in Figure 1D. B. **Stacked bar charts comparing target assay class distributions of all target assays (N=218) to the targets represented in the target-ADR associations (N=51, see also Figure 4B).** No significant differences were observed (χ^2^-test: p-value = 0.09), although the “other” class (grey) had slightly lower representation among the associations. This is expected, since very few drugs were tested on these “other” class assays (Figure 1D), and data availability itself affects the representation in the associations (Figure 4C). Indeed, without considering this “other” class, the distributions were very similar (χ^2^-test: p-value = 0.90), further validating that our algorithm does not bias towards any particular target class.

## Acknowledgements

We are grateful to Mirjam Trame and Andy Stein for giving us the opportunity to participate in the 2018 Novartis Quantitative Sciences Academia-to-Industry Hackathon organized at the Novartis Institutes for Biomedical Research. We also thank Changchang Liu and Xinrui (Sandy) Zou for their contributions to the project at the Hackathon.

## Author contributions

R.I., S.A., A.X.C., S.F., B.K., D.A. and L.U. conceived the study. D.A., A.F. and L.U. provided the Novartis *in vitro* pharmacology data, advice and mentorship. S.A., R.I., A.X.C., S.F., B.K., W.D.M. and J.S. performed data analysis. S.A. developed the random forest modeling. R.I. developed the formalism for target-ADR association inference. S.F., R.I. and A.X.C. developed the query of OpenFDA. J.S. performed the SIDER analysis. S.F., J.S. and R.I. performed the systematic PubMed query. S.A., R.I., and A.X.C. wrote the paper and designed the figures with input from all the authors.

## Conflict of interest

Authors declare no conflict of interest.

## Notes

#### Summary of Updates

Supplemental Figure 1 added. Supplemental data sets provided. Link to Github code repository provided.

https://github.com/samanfrm/ADRtarget

## References

1. Institute of Medicine & Committee on Quality of Health Care in America. To Err Is Human: Building a Safer Health System. (National Academies Press, 2000).

2. Lazarou, J., Pomeranz, B. H. & Corey, P. N. Incidence of adverse drug reactions in hospitalized patients: a meta-analysis of prospective studies. JAMA 279, 1200–1205 (1998).

3. Weiss, A. J., Freeman, W. J., Heslin, K. C. & Barrett, M. L. Adverse Drug Events in US Hospitals, 2010 Versus 2014. HCUP Statistical Brief 234, (2018).

4. Lounkine, E. et al. Large-scale prediction and testing of drug activity on side-effect targets. Nature 486, 361–367 (2012).

5. Bowes, J. et al. Reducing safety-related drug attrition: the use of in vitro pharmacological profiling. Nat. Rev. Drug Discov. 11, 909–922 (2012).

6. Witek, R. P. & Bonzo, J. A. Perspective on In Vitro Liver Toxicity Models. Applied In Vitro Toxicology 4, 229–231 (2018).

7. Whitebread, S., Hamon, J., Bojanic, D. & Urban, L. Keynote review: in vitro safety pharmacology profiling: an essential tool for successful drug development. Drug Discov. Today 10, 1421–1433 (2005).

8. Liu, K. et al. Chemi-net: a graph convolutional network for accurate drug property prediction. arXiv [cs.LG] (2018).

9. Ekins, S. Predicting undesirable drug interactions with promiscuous proteins in silico. Drug Discov. Today 9, 276–285 (2004).

10. Lynch, J. J., 3rd, Van Vleet, T. R., Mittelstadt, S. W. & Blomme, E. A. G. Potential functional and pathological side effects related to off-target pharmacological activity. J. Pharmacol. Toxicol. Methods 87, 108–126 (2017).

11. Hamon, J. et al. In vitro safety pharmacology profiling: what else beyond hERG? Future Med. Chem. 1, 645–665 (2009).

12. Mirams, G. R. et al. Simulation of multiple ion channel block provides improved early prediction of compounds’ clinical torsadogenic risk. Cardiovasc. Res. 91, 53–61 (2011).

13. Chen, A. W. Predicting adverse drug reaction outcomes with machine learning. International Journal Of Community Medicine And Public Health 5, 901–904 (2018).

14. Nguyen, P. A., Born, D. A., Deaton, A. M., Nioi, P. & Ward, L. D. Phenotypes associated with genes encoding drug targets are predictive of clinical trial side effects. bioRxiv 285858 (2018). doi: 10.1101/285858

15. U.S. Food and Drug Administration (FDA). Questions and Answers on FDA’s Adverse Event Reporting System (FAERS). Available at: https://www.fda.gov/drugs/surveillance/fda-adverse-event-reporting-system-faers.

16. Takarabe, M., Kotera, M., Nishimura, Y., Goto, S. & Yamanishi, Y. Drug target prediction using adverse event report systems: a pharmacogenomic approach. Bioinformatics 28, i611–i618 (2012).

17. Mozzicato, P. MedDRA. Pharmaceut. Med. 23, 65–75 (2009).

18. Hauser, A. S., Attwood, M. M., Rask-Andersen, M., Schiöth, H. B. & Gloriam, D. E. Trends in GPCR drug discovery: new agents, targets and indications. Nat. Rev. Drug Discov. 16, 829–842 (2017).

19. for Drug Statistics Methodology, W. C. C. Guidelines for ATC classification and DDD assignment. (2005).

20. U.S. Food and Drug Administration (FDA). openFDA. Available at: https://open.fda.gov.

21. Maciejewski, M. et al. Reverse translation of adverse event reports paves the way for derisking preclinical off-targets. Elife 6, (2017).

22. Dumouchel, W. Bayesian Data Mining in Large Frequency Tables, with an Application to the FDA Spontaneous Reporting System. Am. Stat. 53, 177–190 (1999).

23. Fram, D. M., Almenoff, J. S. & DuMouchel, W. Empirical Bayesian Data Mining for Discovering Patterns in Post-marketing Drug Safety. in Proceedings of the Ninth ACM SIGKDD International Conference on Knowledge Discovery and Data Mining 359–368 (ACM, 2003).

24. Almenoff, J. S., LaCroix, K. K., Yuen, N. A., Fram, D. & DuMouchel, W. Comparative Performance of Two Quantitative Safety Signalling Methods. Drug Saf. 29, 875–887 (2006).

25. DuMouchel, W. & Harpaz, R. Regression-adjusted GPS algorithm (RGPS) April 2015. Oracle Health Sci (2015).

26. Sequeira, A. et al. Alpha 2A adrenergic receptor gene and suicide. Psychiatry Res. 125, 87–93 (2004).

27. Cottingham, C. & Wang, Q. α2 adrenergic receptor dysregulation in depressive disorders: Implications for the neurobiology of depression and antidepressant therapy. Neurosci. Biobehav. Rev. 36, 2214–2225 (2012).

28. Kuhn, M., Letunic, I., Jensen, L. J. & Bork, P. The SIDER database of drugs and side effects. Nucleic Acids Res. 44, D1075–9 (2016).

29. Curran, M. E. et al. A molecular basis for cardiac arrhythmia: HERG mutations cause long QT syndrome. Cell 80, 795–803 (1995).

30. Wen, D., Liu, A., Chen, F., Yang, J. & Dai, R. Validation of visualized transgenic zebrafish as a high throughput model to assay bradycardia related cardio toxicity risk candidates. J. Appl. Toxicol. 32, 834–842 (2012).

31. Mitcheson, J. S. hERG potassium channels and the structural basis of drug-induced arrhythmias. Chem. Res. Toxicol. 21, 1005–1010 (2008).

32. Yusof, I., Shah, F., Hashimoto, T., Segall, M. D. & Greene, N. Finding the rules for successful drug optimisation. Drug Discov. Today 19, 680–687 (2014).

33. Movsesian, M., Wever-Pinzon, O. & Vandeput, F. PDE3 inhibition in dilated cardiomyopathy. Curr. Opin. Pharmacol. 11, 707–713 (2011).

34. Knight, W. & Yan, C. Therapeutic potential of PDE modulation in treating heart disease. Future Med. Chem. 5, 1607–1620 (2013).

35. Ely, S. W., Matherne, G. P., Coleman, S. D. & Berne, R. M. Inhibition of adenosine metabolism increases myocardial interstitial adenosine concentrations and coronary flow. J. Mol. Cell. Cardiol. 24, 1321–1332 (1992).

36. Aviña-Zubieta, J. A. et al. Immediate and past cumulative effects of oral glucocorticoids on the risk of acute myocardial infarction in rheumatoid arthritis: a population-based study. Rheumatology 52, 68–75 (2013).

37. Oakley, R. H. & Cidlowski, J. A. Glucocorticoid signaling in the heart: A cardiomyocyte perspective. J. Steroid Biochem. Mol. Biol. 153, 27–34 (2015).

38. Huerta, C., Castellsague, J., Varas-Lorenzo, C. & García Rodríguez, L. A. Nonsteroidal anti-inflammatory drugs and risk of ARF in the general population. Am. J. Kidney Dis. 45, 531–539 (2005).

39. Wang, L. et al. Podocyte-specific knockout of cyclooxygenase 2 exacerbates diabetic kidney disease. Am. J. Physiol. Renal Physiol. 313, F430–F439 (2017).

40. Slattery, P., Frölich, S., Schreiber, Y. & Nüsing, R. M. COX-2 gene dosage-dependent defects in kidney development. Am. J. Physiol. Renal Physiol. 310, F1113–22 (2016).

41. Ye, H. et al. Modulation of Polycystic Kidney Disease Severity by Phosphodiesterase 1 and 3 Subfamilies. J. Am. Soc. Nephrol. 27, 1312–1320 (2016).

42. Wadhwa, S., Wadhwa, P., Dinda, A. K. & Gupta, N. P. Differential expression of potassium ion channels in human renal cell carcinoma. Int. Urol. Nephrol. 41, 251–257 (2009).

43. Fagerberg, L. et al. Analysis of the human tissue-specific expression by genome-wide integration of transcriptomics and antibody-based proteomics. Mol. Cell. Proteomics 13, 397–406 (2014).

44. Nussey, S. S. & Whitehead, S. A. Endocrinology: an integrated approach. (CRC Press, 2013).

45. Miyamoto, J. et al. The pituitary function of androgen receptor constitutes a glucocorticoid production circuit. Mol. Cell. Biol. 27, 4807–4814 (2007).

46. Nuzzi, R., Scalabrin, S., Becco, A. & Panzica, G. Gonadal Hormones and Retinal Disorders: A Review. Front. Endocrinol. 9, 66 (2018).

47. Ashton, H. Guidelines for the Rational Use of Benzodiazepines. Drugs 48, 25–40 (1994).

48. Kubitz, R., Dröge, C., Stindt, J., Weissenberger, K. & Häussinger, D. The bile salt export pump (BSEP) in health and disease. Clin. Res. Hepatol. Gastroenterol. 36, 536–553 (2012).

49. Riede, J., Poller, B., Huwyler, J. & Camenisch, G. Assessing the Risk of Drug-Induced Cholestasis Using Unbound Intrahepatic Concentrations. Drug Metab. Dispos. 45, 523–531 (2017).

50. Liu, L.-Y., Wang, X.-H., Lu, Y., Zhu, Q.-R. & Wang, J.-S. Association of variants of ABCB11 with transient neonatal cholestasis: ABCB11 and TNC. Pediatr. Int. 55, 138–144 (2013).

51. Geenes, V. & Williamson, C. Intrahepatic cholestasis of pregnancy. World J. Gastroenterol. 15, 2049 (2009).

52. Liu, M. et al. Large-scale prediction of adverse drug reactions using chemical, biological, and phenotypic properties of drugs. J. Am. Med. Inform. Assoc. 19, e28–35 (2012).

53. Bender, A. et al. Analysis of pharmacology data and the prediction of adverse drug reactions and off-target effects from chemical structure. ChemMedChem: Chemistry Enabling Drug Discovery 2, 861–873 (2007).

54. Dawson, S., Stahl, S., Paul, N., Barber, J. & Kenna, J. G. In vitro inhibition of the bile salt export pump correlates with risk of cholestatic drug-induced liver injury in humans. Drug Metab. Dispos. 40, 130–138 (2012).

55. Montanari, F. et al. Flagging Drugs That Inhibit the Bile Salt Export Pump. Mol. Pharm. 13, 163–171 (2016).

56. Cheng, J. & Grande, J. P. Cyclic nucleotide phosphodiesterase (PDE) inhibitors: novel therapeutic agents for progressive renal disease. Exp. Biol. Med. 232, 38–51 (2007).

57. Tatonetti, N. P., Ye, P. P., Daneshjou, R. & Altman, R. B. Data-driven prediction of drug effects and interactions. Sci. Transl. Med. 4, 125ra31 (2012).

58. Breiman, L. Random Forests. Mach. Learn. 45, 5–32 (2001).

59. Charte, F. & Charte, D. Working with Multilabel Datasets in R: The mldr Package. R J. 7, (2015).

60. Rivolli, A. & de Carvalho, A. C. The utiml Package: Multi-label Classification in R. R J. (2018).

61. Saito, T. & Rehmsmeier, M. Precrec: fast and accurate precision--recall and ROC curve calculations in R. Bioinformatics 33, 145–147 (2017).

62. Menze, B. H. et al. A comparison of random forest and its Gini importance with standard chemometric methods for the feature selection and classification of spectral data. BMC Bioinformatics 10, 213 (2009).

63. Brugada, R. et al. Sudden death associated with short-QT syndrome linked to mutations in HERG. Circulation 109, 30–35 (2004).

64. Pinto, C. S. et al. Phosphodiesterase Isoform Regulation of Cell Proliferation and Fluid Secretion in Autosomal Dominant Polycystic Kidney Disease. J. Am. Soc. Nephrol. 27, 1124–1134 (2016).

65. Wang, X., Ward, C. J., Harris, P. C. & Torres, V. E. Cyclic nucleotide signaling in polycystic kidney disease. Kidney Int. 77, 129–140 (2010).

66. Babcock, J. J. & Li, M. hERG channel function: beyond long QT. Acta Pharmacol. Sin. 34, 329–335 (2013).

67. Mertens, K. L., Kalsbeek, A., Soeters, M. R. & Eggink, H. M. Bile Acid Signaling Pathways from the Enterohepatic Circulation to the Central Nervous System. Front. Neurosci. 11, 617 (2017).

68. Srivastava, A. Progressive familial intrahepatic cholestasis. J. Clin. Exp. Hepatol. 4, 25–36 (2014).

69. Strautnieks, S. S. et al. A gene encoding a liver-specific ABC transporter is mutated in progressive familial intrahepatic cholestasis. Nat. Genet. 20, 233–238 (1998).

70. Watanabe, M. et al. Bile acids induce energy expenditure by promoting intracellular thyroid hormone activation. Nature 439, 484–489 (2006).

71. Mukaisho, K.-I. et al. High serum bile acids cause hyperthyroidism and goiter. Dig. Dis. Sci. 53, 1411–1416 (2008).

72. Comhair, S. A. A. et al. Metabolomic Endotype of Asthma. J. Immunol. 195, 643–650 (2015).

73. Tan, X. et al. Genetic and Proteomic characterization of Bile Salt Export Pump (BSEP) in Snake Liver. Sci. Rep. 7, 43556 (2017).

74. Li, C.-H., Sung, F.-C., Wang, Y.-C., Lin, D. & Kao, C.-H. Gallstones increase the risk of developing renal stones: a nationwide population-based retrospective cohort study. QJM 107, 451–457 (2014).

75. Li, C.-H., Sung, F.-C., Wang, Y.-C., Lin, D. & Kao, C.-H. Gallstones increase the risk of developing renal stones: a nationwide population-based retrospective cohort study. QJM 107, 451–457 (2014).

